# Divergent spontaneous antibiotic-resistance evolution confers reciprocal and exploitable collateral sensitivity effects

**DOI:** 10.1101/2025.09.22.677096

**Authors:** Apostolos Liakopoulos, Sebastian T. Tandar, Irene V. Hoogendijk, Efthymia Fragkiskou, Zheren Zhang, Bastienne Vriesendorp, Linda B. S. Aulin, J. G. Coen van Hasselt, Daniel E. Rozen

## Abstract

The global rise of antibiotic-resistant pathogens has outpaced the development of new antibiotics, prompting the urgent need for alternative treatment strategies. One such approach is to leverage collateral sensitivity (CS), where resistance to one antibiotic increases susceptibility to another. However, the clinical implementation of CS-based therapies depends on the consistency of these responses, which is challenged by variable resistance mutations and the dynamics of resistant strains during infection. Here, we combined experiments and mathematical models to assess the consistency and consequences of CS responses in the Gram-positive pathogen *Streptococcus pneumoniae* following the *de novo* acquisition of resistance to five commonly used antibiotics. We found that many collateral responses were unpredictable and inconsistent between different resistance mutations. However, for two antibiotic pairs, we identified consistent unidirectional (RIF → FUS) and bidirectional (LNZ ↔ FUS) CS interactions, despite the divergent evolutionary trajectories of resistant strains, as revealed by whole-genome sequencing. To evaluate if CS for these combinations can be exploited to design dosing strategies to eradicate *S. pneumoniae* infections while suppressing resistance, we developed a mathematical stochastic pharmacokinetic-pharmacodynamic (PK-PD) model, which integrated our experimentally derived PD parameters with existing clinical PK models. Our model-based analyses confirmed the superiority of these antibiotic combinations over monotherapy and showed that their efficacy depends on the presence of CS interactions between the administered antibiotics. In summary, our study demonstrates how consistent CS interactions can be leveraged to inform treatment strategies, laying the groundwork for CS-guided therapies to preserve antibiotic efficacy.

## Introduction

The widespread use of antimicrobial agents in medicine and agriculture has driven the global rise of drug-resistant pathogens and poses a major challenge to public health^1,2^. Among the most concerning threats is *Streptococcus pneumoniae* which remains a leading cause of community-acquired pneumonia, meningitis, and bacteremia^3^. Single- and multidrug-resistant *S. pneumoniae* strains are becoming increasingly common against a broad variety of antibiotics^4,5^, thereby limiting treatment options and affecting disease burden. Antibiotic-resistant strains have historically been managed through the introduction of a new antibiotic class or treatment regime. However, because of significant declines in new antibiotic development, alternative strategies are urgently needed.

Antibiotic combination therapy has emerged as a promising alternative to counteract resistance. Antibiotic combinations are often selected based on their synergistic effects that can enhance the rate and efficacy of bacterial clearance^6,7^. However, while this approach can be effective in rapidly eliminating infections, it may also impose strong selective pressures on bacterial populations that can promote the invasion and persistence of antibiotic-resistant strains^8^. Another approach to antibiotic combination therapy involves leveraging collateral effects, a phenomenon in which resistance to one antibiotic affects susceptibility to a second antibiotic^9^. A collateral effect that increases sensitivity to a second antibiotic is known as collateral sensitivity (CS) and an effect that decreases sensitivity as collateral resistance (CR). While the occurrence of CR is regarded as a negative outcome for combination therapies, CS provides a vulnerability that can be exploited during antibiotic treatment to reduce the risks of resistance^10–12^. According to CS-based strategies, cycling or mixing antibiotics effectively penalizes bacteria that develop resistance to one antibiotic by exposing them to another to which they have become more sensitive. By systematically alternating or co-administering antibiotics based on their CS relationships, this approach can help counteract resistance while maintaining treatment efficacy^11,13^.

An important prerequisite of CS-based strategies is that collateral responses are conserved in strains carrying different mutations that confer resistance to the same antibiotic^14–16^. Conservation is essential to ensure that collateral responses to pairs of antibiotics are predictable and reliable, even if resistance is driven by different mutations. Screens to assess the predictability and conservation of collateral responses have produced mixed results. While CS is highly predictable for some antibiotic pairs in *Escherichia coli*^17^ and *Pseudomonas aeruginosa*^15,16^, responses in most cases are less consistent. In our previous work in *S. pneumoniae*, for example, we found that different mutations in *gyrA* or *parC* that confer resistance to fluoroquinolones led to consistent CS responses against 7 of 12 tested antibiotics. However, for the remaining 5 antibiotics, the same mutations either had no collateral effects at all, or even led to a mixture of CS and CR^12^. While such screens help to identify potentially (un)suitable antibiotic pairs for CS-based therapies, we lack a comprehensive investigation of CS across different antibiotics in *S. pneumoniae*, thus hampering the ability to use this strategy to establish treatment protocols. In addition, previous studies have not accounted for the potential impact of the within-strain population diversity caused by the coexistence of multiple resistant genotypes during infection. These genotypes, arising through different spontaneous mutations, can lead to heterogeneous bacterial populations with variable levels of antibiotic resistance, collateral responses, as well as growth characteristics^18,19^. Thus, applying CS-based strategies in the clinic requires assessing both the consistency of CS development and the population dynamics of coexisting resistant strains.

The aim of this study was to determine the extent to which CS can be exploited in treatment strategies against *S. pneumoniae*. We combined experiments and simulations to investigate the emergence, repeatability and magnitude of CS responses against a panel of clinically relevant antibiotics. For selected CS pairs, we conducted static time-kill experiments to further characterize the pharmacodynamics (PD) of the antibiotics against *S. pneumoniae*. Finally, to assess the clinical applicability of the identified CS pairs to suppress infection and prevent the establishment of resistant bacterial population, we established and used a stochastic pharmacokinetics-pharmacodynamics (PK-PD) model to predict the outcomes of antibiotic combination treatments. The outcome of this combined experimental and modelling study offers valuable insights into the use of CS-based antibiotic combinations for managing antibiotic-resistant *S. pneumoniae* infections, while also providing a strategy to preserve the effectiveness and extend the lifespan of last-resort antibiotics.

## Methods

### Bacterial strains and growth conditions

All experiments were performed with the *S. pneumoniae* R6 strain^20^. Ten independent spontaneously resistant mutants were isolated for each of five antibiotics, as described below. *S. pneumoniae* ATCC 49619^21^ was used as a quality control for antimicrobial susceptibility testing. Strains used in this study are listed in **Table S1**. Unless otherwise mentioned, all strains were grown at 37°C under 5% CO2 for 18-24 hours either on tryptic soy agar for solid culture (TSYA) (BD, New Jersey, USA) supplemented with 0.5% w/v yeast extract (BD, New Jersey, USA) and 5% v/v sheep blood (Sanbio, Uden, The Netherlands), or in tryptic soy broth supplemented with 0.5% w/v yeast extract and 5% v/v sheep blood (TSYB) for liquid culture. The 13 antibiotics used in this study were prepared from powder stock and stored at -20°C or -80°C according to the manufacturers’ recommendations (**Table S2**).

### Selection of antibiotic-resistant mutants

Resistant mutants were selected against five antibiotics with different modes of action to which resistance in *S. pneumoniae* can arise *via* chromosomal mutations^1^: ciprofloxacin (CIP), fusidic acid (FUS), linezolid (LZD), rifampicin (RIF) or trimethoprim/sulfamethoxazole (SXT). Selection of drug-resistant isolates for each antibiotic was performed using a modified fluctuation-test approach by inoculating approximately 10^3^ colony-forming units (CFU) into ten independent tubes containing 10 ml TSYB and growing without agitation to an optical density at 620 nm (OD_620_) of 0.3, corresponding to ∼2 × 10^8^cells/ml. After incubation, cells were collected from each culture by centrifugation (10 minutes; 5000 rpm) and then resuspended in 100 μl 0.85% w/v saline. Resuspended pellets were plated on a single agar plate containing each antibiotic at a concentration equal to the epidemiological cut-off value (ECOFF)^22^. To isolate independent mutants, we randomly selected a single colony per plate from the 10 different plates for each antibiotic, for a total of 50 strains (five antibiotics; 10 strains each). Species identity of all antibiotic-resistant mutants was confirmed using species-specific amplification of *lytA*^23^ and Matrix-Assisted Laser Desorption/Ionization with Time-of-Flight mass spectrometry (MALDI-TOF; Bruker, Massachusetts, USA) according to manufacturer’s recommendations.

### Whole-genome sequencing

Genomic DNA was extracted from the *S. pneumoniae* R6 WT and all 50 antibiotic-resistant mutants using the DNeasy Blood & Tissue Kit (QIAGEN, Hilden, Germany) according to the manufacturer’s recommendations for Gram-positive bacteria. Quality control of purified genomic DNA was performed using Quant-iT™ dsDNA BR Assay Kit (ThermoFisher Scientific, Breda, The Netherlands) and 1% agarose gel electrophoresis. Genomic DNA was fragmented by ultrasound on a Covaris S/E210 ultrasonicator (Covaris, Brighton, UK) and was subsequently used for library preparation^24^. Whole genome sequencing was performed using 100-bp paired-end libraries on a BGISEQ-500 platform at BGI Europe Genome Center (Copenhagen, Denmark). High-quality filtered reads were mapped to the reference genome of *S. pneumoniae* D39V available in GenBank (accession number NZ_CP027540.1) using the open-source computational pipeline *breseq*^25^ with the default parameter settings in order to identify and annotate genetic differences found between our resistant mutants and the reference R6 WT strain.

### Antibiotic susceptibility assays and collateral effect determination

Minimum inhibitory concentrations (MICs) of all selected resistant strains were determined in triplicate by broth microdilution^26^, using a 1.5-fold testing scale to include the standard two-fold antibiotic concentrations and their median values. Mueller-Hinton cation-adjusted broth (MHCAB; BD, New Jersey, USA) was supplemented with 100 U/ml of catalase (Worthington Biochemical Corporation, New Jersey, USA) in place of 5% lysed horse blood to detoxify *S. pneumoniae*-derived hydrogen peroxide under aerobic conditions, which otherwise slows growth and can trigger autolysis, as previously described^27^.

Collateral effects exhibited by a resistant mutant strain M (*CE*_*M*_) towards an antibiotic were quantified as the log2-scaled fold change in the MICs between each mutant and the parental R6 WT strain (**Eq. 1**).

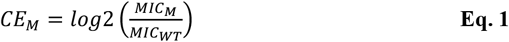

In the above equation, MIC_M_ and MIC_WT_ represent the MIC of the mutant and the parental R6 WT strain, respectively. Collateral effects were categorized as CS when *CE*_*M*_ was negative (indicating a decrease in MIC), and CR when *CE*_*M*_ was positive (indicating an increase in MIC). The conservation of these effects was evaluated by assessing the consistency of the collateral response (CS or CR) across strains resistant to the tested antibiotics, using CS_50_ and CR_50_ thresholds. These thresholds were defined as CS or CR effects occurring in more than 50% of the mutants tested^12^.

### Partial least squares regression for the quantification of mutated loci impact on antibiotic sensitivity

Partial least squares (PLS) regression was used to evaluate the impact of mutated loci on shifts in MIC for specific antibiotics in mutant strains. In this analysis, the presence or absence of a mutation in a given locus (coded as 1 or 0, respectively) was used as the independent variable, while the log_2_ MIC shift from the parental strain served as the dependent variable. PLS was performed using 1 to 20 principal components. The final number of components was selected based on the root mean squared error of prediction (RMSEP), choosing the smallest number of components that achieved at least a 95% reduction in RMSEP from its maximum value (with one component) to the minimum observed across all tested components.

The resulting PLS coefficients represent the magnitude and direction of the impact of a given mutated locus on the log_2_ MIC shift. Positive coefficient values indicate a contribution to increased resistance (higher MIC), while negative values indicate increased sensitivity. Larger absolute values reflect a stronger influence on the MIC shift. PLS was performed in R using the package ‘*pls*’.

### Static time-kill assays

Static time-kill assays were performed on *S. pneumoniae* R6 WT to characterize the PD of the antibiotics exhibiting conserved and bidirectional CS effects, namely FUS, LNZ, and RIF. Assays were performed in triplicate with a starting inoculum of ∼ 1.5 × 10^6^ CFU/mL in 15 mL of MHCAB supplemented with 5% lysed blood. FUS, LNZ, and RIF were added to the culture to reach a final concentration of 0X, 0.5X, 1X, 2X, 4X, 16X or 32X its respective MIC of the parental strain (**Table S3**). The resulting liquid cultures were incubated under continuous agitation (80 rpm). Samples (100 µL) were collected at 0, 1, 2, 4 and 6 h post inoculation. The collected samples were serially diluted and plated on TSYA agar plates for viable cell titer determination after 48 hours incubation.

As stationary phase was not reached within the 6 hours of the time-kill assays, the PD of FUS, LNZ, and RIF was characterized from the assay without assuming population limitation (**Eq. 2**). The concentration-effect relationship of FUS, LNZ, and RIF towards the bacterial population were modelled using the Hill equation, defined by the parameters *v*_*death,max*_ (maximum bactericidal effect), *EC*_50_ (concentration to achieve 50% of the bactericidal effect), and *h* (Hill exponent of the concentration-effect curve). These parameters were estimated based on observations from the time-kill assay by fitting **Eq. 2** to the observation values on a log-linear scale.

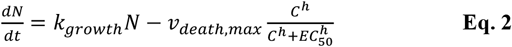

### Checkerboard assays

Antibiotic interactions between the antibiotics exhibiting conserved and bidirectional CS effects (FUS, LNZ, and RIF) were assessed in triplicate by checkerboard assays on the *S. pneumoniae* R6 WT strain. The checkerboard assay measures bacterial growth inhibition across a matrix of concentration combinations for two antibiotics. In this study, checkerboard assays were used to assess interactions between FUS-LNZ as well as FUS-RIF. Checkerboard assays were carried out using a starting inoculum of ∼ 1.5 × 10^6^ CFU/mL in MHCAB (BD, New Jersey, USA) supplemented with 100 U of catalase (Worthington Biochemical Corporation, New Jersey, USA). The ranges of FUS, LNZ, and RIF concentrations tested were 0.25 - 256, 0.03 - 32, and 0.00006 - 0.0625 mg/L, respectively. The generated data were analysed to identify antibiotic interactions using the FIC index (FICI) and interpreted according to previously established criteria, in which interactions were categorized as synergistic (FICI ≤ 0.5), antagonistic (FICI > 4.0), or no interaction (FICI > 0.5-4.0)^28^.

### Growth rate measurements

Growth rates were determined in triplicate for *S. pneumoniae* R6 WT and the FUS-, LNZ-, and RIF-resistant mutants. For this, an initial inoculum of each strain was prepared by transferring colonies grown on TSYA to tubes of TSYB to achieve an OD_620_ of 0.3, corresponding to ∼2 × 10^8^ cells/ml. Then, 50 µl of this inoculum was transferred to 5 ml of TSYB pre-culture and incubated without agitation for 18-24 hours. On the following day, 20 µl each pre-culture was used to inoculate 180 µl of TYSB medium in a 96-well plate (transparent, Greiner Bio-One). Inoculated assay plates were placed in an incubated automated plate reader (lower limit of quantification; LoQ of 0.05 a. u.; BioTek Synergy HT, Agilent). OD_620_ was measured every 30 minutes for the next 24 hours with 20 seconds of agitation before each measurement. OD_620_ measurements were used to describe the growth kinetics of each strain *i* and derive its growth rate (*k’*_*growth*_) and maximum cell density (*N’*_*max*_) using a population-limited growth model (**Eq. 3**).

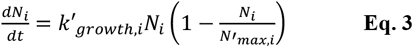

An empirical delay term ***t***_***delay***_ was included to account for initial growth where cell density was below the detection limit of the plate reader, as well as growth delay that may occur when the bacteria were adjusting to the new growth environment after inoculation. In this model, the change in bacterial population density was set to zero until ***t***_***delay***_ had elapsed. A random effect was applied to the inoculum size *N*_*0*_ to capture inter-replicate variability in inoculum size. Model fitting was performed using the package ‘*nlme’* in R by implementing inter-replicate variability on inoculum size *N*_*0*_.

Growth rates measured in microtiter plates may differ from those in static time-kill experiments due to differences in culture conditions (*e*.*g*., volume, vessel size) and measurement methods. The CFU-based growth rate of each mutant strain *i* (*k*_*growth,i*_) was approximated from its OD-based growth rate (*k*^′^_*growth,i*_), assuming that the ratio between CFU- and OD-based growth rates is constant across all strains (**Eq. 4**).

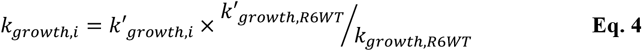

### Pharmacokinetic-pharmacodynamic model development

A mathematical PK-PD model was developed to approximate the potential outcome of antibiotic treatment in an infection caused by *S. pneumoniae*. To this end, PK-PD models were implemented, incorporating the stochastic emergence of (multiple) resistant bacterial sub-populations whose phenotypes were based upon our experimental measurements. These stochastic PK-PD models were developed by integrating published PK models with the PD model developed in this study (**Fig. 1**). Published population PK models for FUS^29^, LNZ^30^, and RIF^31^ were used to simulate antibiotic treatments for the respective antibiotics. Unbound plasma concentrations were calculated based on reported plasma protein binding ratios for FUS (97.7%)^32^, LNZ (17.7%)^30^, and RIF (88.9%)^33^. To standardize simulations, individual patient characteristics required for the PK models were set to the median or standard values reported in the original studies (**Table S4**). Model simulations were performed to estimate the median concentration–time profiles of FUS, LNZ, and/or RIF during specified treatment regimens. The PD model described bacterial growth using a multi-population model, including one (homogeneous) population of antibiotic-sensitive *S. pneumoniae* cells representing the R6 WT strain as well as two heterogeneous populations of mutant cells, each resistant to a specific antibiotic. Each resistant population was further divided into 10 sub-populations of cells, corresponding to the 10 mutants per-antibiotic generated in this study. *De novo* resistance mutations were assumed to occur stochastically and unidirectionally from the R6 WT strain to one of the resistant sub-populations, with each event occurring at equal probability. A complete description of the PD model is provided in **Supplementary Material 2**.

**Fig. 1.**
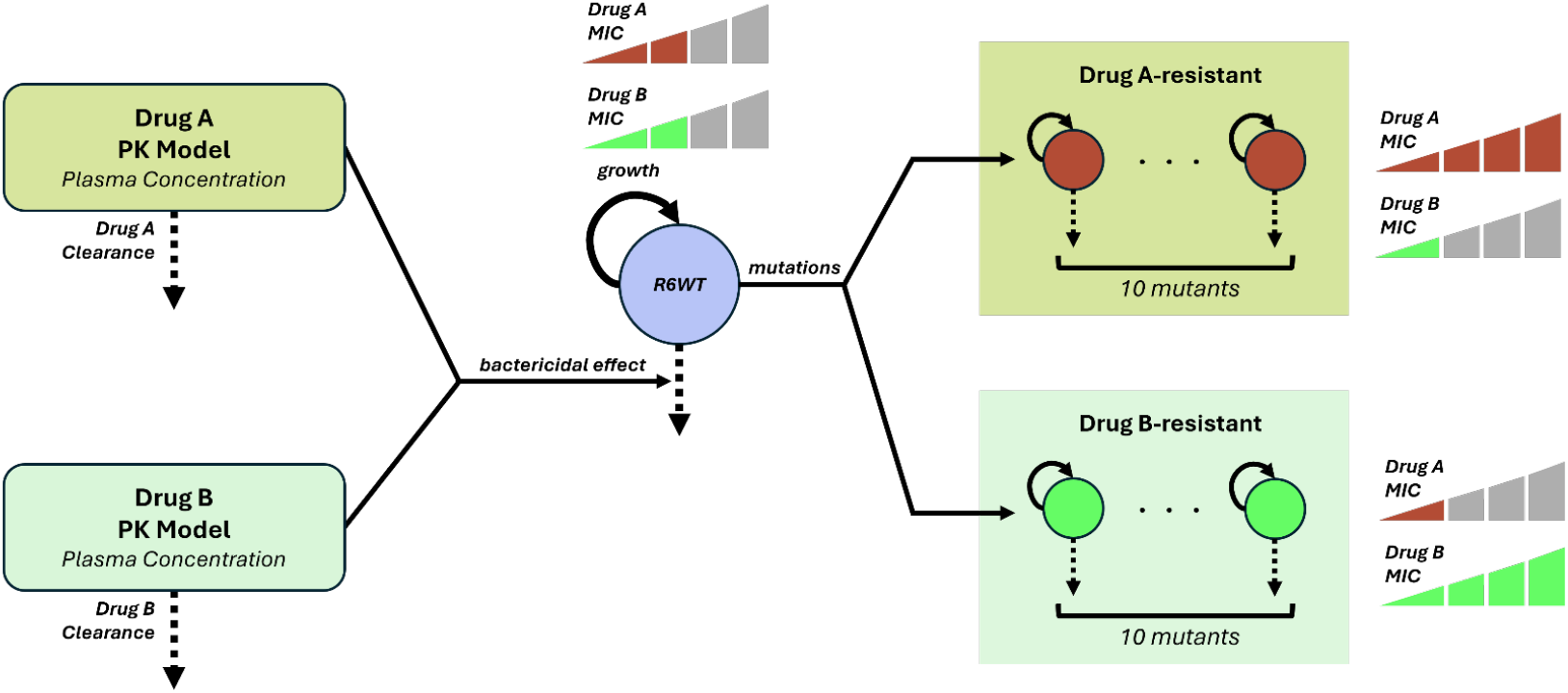
Schematic diagram of the PK-PD model structure. The PK-PD models simulate the concentration–time profiles and the corresponding bacterial responses to two antibiotics (indicated as drug A and drug B with bidirectional collateral sensitivity between them in the example depicted) used in combination. Published population PK models were used to estimate the median plasma concentrations of each antibiotic. Antibiotic concentrations exert varying levels of bactericidal activity on each bacterial (sub-) population. The model consists of one wildtype R6 WT bacterial population and two mutant populations, each resistant to either drug A or drug B; each mutant population is further divided into 10 bacterial sub-populations to represent the 10 mutant strains characterized in this study for each antibiotic. Each bacterial (sub-)population grows at its own specific growth rate. Mutations from the R6 WT strain were assumed to occur at equal rates to all mutant sub-populations.

### *In silico* evaluation of antibiotic treatments

Our stochastic PK-PD model was used to simulate the potential outcome of antibiotic treatment against *S. pneumoniae* bacteraemia. A total blood volume of 5 L (average adult plasma volume)^34^ was assumed for all simulations, representing a hypothetical treatment of a bacteraemia infection. To replicate the natural occurrence of mutants during a *S. pneumoniae* infection, an infection with a starting population size of 10 CFU/mL was allowed to reach a cell density of 10^4^ CFU/mL (established *S. pneumoniae* bacteraemia^35^) before treatment was initiated. Antibiotic treatment was assumed to last for a period of 14 days. A final cell density of < 10^3^ CFU/mL was used as the target of therapeutic efficacy. The stochastic simulation was replicated 500 times. The fraction of these 500 replicate simulations that reached this threshold cell density was used as a measure of treatment efficacy. The standard clinical doses for FUS, RIF, and LNZ used in The Netherlands^36^ (**Table S5**) were used for the simulations. Each simulation included 21 bacterial sub-populations: the R6 WT parental strain plus 20 sub-populations resistant to two different drugs. In combination therapy simulations, 10 sub-populations were resistant to each of the two drugs in the combination. In monotherapy simulations, 10 sub-populations were resistant to the treatment drug, and 10 were resistant to another drug, chosen arbitrarily from the remaining two. This design ensured that the overall rate of resistance was consistent across all treatment scenarios.

To evaluate the applicability of an antibiotic treatment combination using a CS antibiotic pair (drug A and drug B), three possible regimens were evaluated for each drug combination. These included 1) co-administration (mixing) of drug A and drug B, or 12-hours cycling starting with 2) drug A or 3) drug B. The dose of each drug in the combination was either kept constant at its equivalent daily standard dose, or half the standard amount when 12-hours cycling was used.

To assess the importance of the CS phenotype during combination treatment, parallel simulations on each of the three dosing regimens were performed where collateral antibiotic effects (both CS and CR) were excluded. This was done by fixing the MIC of the mutant strains for antibiotics they were not selected for to match the MIC of the parental R6 WT strain.

### Simulation software

Deterministic simulation using the population pharmacokinetic model was performed using the ‘*rxode2*’ package in R. Stochastic simulation of bacterial density using the PK-PD model was performed on R using the τ-leap Gillespie method^37^ (τ fixed to 0.1 hours) and by applying stochasticity on mutational events. The expected occurrence of mutations into each mutant sub-population at each timestep *τ* was assumed to be binomially distributed. The stochastic simulation was replicated 500 times for each treatment condition.

## Results

### Spontaneous antibiotic-resistance evolution confers predominantly collateral sensitivity

To systematically study collateral and fitness effects associated with the development of antibiotic resistance in *S. pneumoniae*, we isolated 50 spontaneously evolved strains with resistance against one of five antibiotics with different modes of action (CIP, FUS, LNZ, RIF and SXT). In total, we selected 10 independent mutants per antibiotic. The recovered mutants showed increased MICs to the antibiotics they were selected against (**Fig. S1**; parental MICs shown on **Table S3**), and most showed comparable growth to the parental R6 WT strain (**Fig. S2** and **S3, Table S6**). The exceptions were mutant RIF9 and and LNZ5, which exhibited statistically significant (p < 0.05) increases in growth rate and maximum cell density, respectively (**Fig. S3**). Collateral effects in the recovered antibiotic-resistant mutants were assessed against 13 antibiotics. This included eight antibiotics commonly used to treat Gram-positive pathogens and the five antibiotics that were used to select for the resistant strains (**Table S2**). This resulted in 60 unique antibiotic pairs, and 600 antibiotic pair-strain sets evaluated in the analysis. Collateral effects were observed in approximately 60% (362/600) of all tested antibiotic pair-strain sets, with CS identified in about 42% of antibiotic pair-strain sets (**Fig. 2**).

**Fig. 2.**
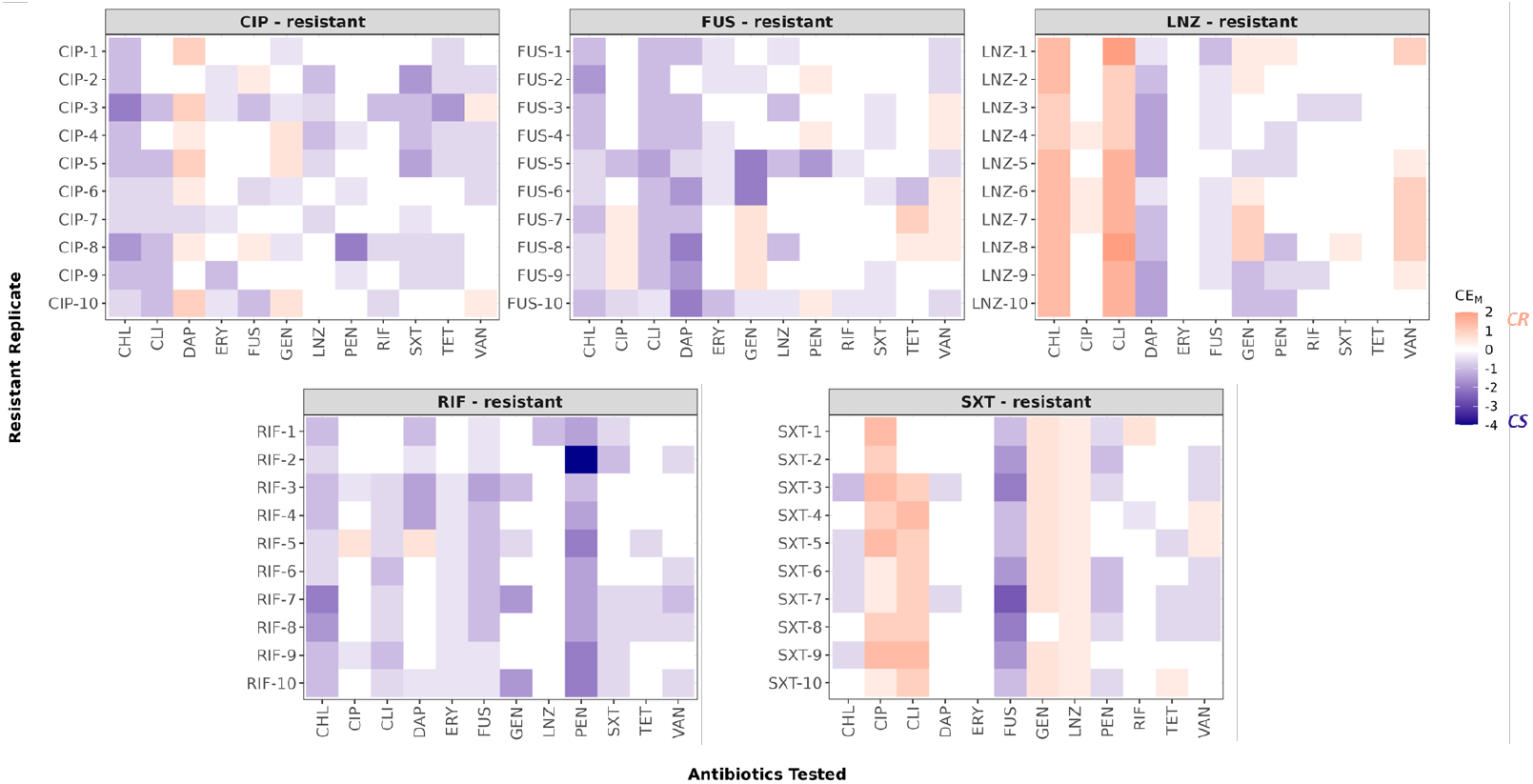
Collateral effect profile of antibiotic-resistant *S. pneumoniae* mutants. For each tested antibiotics, collateral effect magnitude for each corresponding mutant strain (*CE*_*M*_) was calculated as the log2-fold MIC change from that observed for the parental R6 WT strain. A positive value of *CE*_*M*_ denoted CR and was indicated by the colour red. Conversely, a negative *CE*_*M*_ indicated CS interaction and was indicated by blue. Colour intensity indicates the magnitude of the respective collateral effect. CHL, chloramphenicol; CIP, ciprofloxacin; CLI, clindamycin; DAP, daptomycin; ERY, erythromycin; FUS, fusidic acid; GEN, gentamicin; LNZ, linezolid; PEN, penicillin; RIF, rifampicin; SXT, trimethoprim/sulfamethoxazole; TET, tetracycline; VAN, vancomycin.

The observed collateral effects among the independent mutants, even within the same resistant group, varied in number, intensity, and/or direction for each of the 12 antibiotics, with all 50 mutants exhibiting distinct collateral effect pattern from one another. Variability was also evident between resistance groups (CIP, FUS, LNZ, RIF and SXT), with each group of resistant mutants displaying specific and distinct collateral effects to the other antibiotics (**Fig. 2**), thereby highlighting numerous (un)suitable potential combinations for CS-based therapies. CIP-resistant mutants showed CS to several antibiotics, most notably CHL, while also demonstrating conserved CR towards daptomycin (DAP). FUS-resistant mutants predominantly exhibited CS towards CHL and clindamycin (CLI), and somewhat conserved CR towards vancomycin (VAN). LNZ-resistant mutants were characterized by high level CR to CHL and CLI but retained CS to DAP. RIF-resistant mutants displayed broad CS towards a range of antibiotics without any conserved CR, making RIF particularly promising for CS-informed therapies. SXT-resistant mutants demonstrated conserved CS to several antibiotics, including FUS and CHL, but also showed consistent CR to CIP and LNZ. Overall, the patterns of CS and CR varied by resistance mutant and resistant group, but conserved yet infrequent CS trends within each group may be used to inform antibiotic cycling or mixing combination strategies to limit the emergence of multidrug resistance.

### Distinct mutational profiles within and between resistance groups

To identify the genetic changes associated with spontaneous evolution of antibiotic resistance of our mutants, we performed whole-genome sequencing of the recovered mutants and the parental R6 WT strain. Most of the observed resistance phenotypes were associated with genetic changes in chromosomal loci previously implicated in resistance to these antibiotics (primary resistance genes) (**Fig. S4**). In CIP-resistant strains, mutations were predominantly found in *parC* (8 out of 10 isolates)^38^. FUS resistance was consistently associated with mutations in *fusA*^39^, RIF resistance with mutations in *rpoB*^40^, and SXT resistance primarily with mutations in *folA* ^41^ in 8 out of 10 isolates. Notably, within each of these resistance groups, the mutations occurred consistently in the same primary resistance locus, although the specific mutations varied among resistant strains.

In contrast, LNZ resistance was associated with a more diverse mutational landscape. Only 4 out of 10 LNZ-resistant strains carried mutations in loci previously associated with LNZ resistance (*e*.*g*., 23S rRNA, ribosomal proteins, or ABC transporters^42^). The remaining isolates harboured mutations in different genes that, although not previously linked to LNZ resistance, encode proteins with functions analogous to known primary LNZ resistance genes, such as ABC transporters and RNA-methyltransferases^42,43^. Alongside these mutations, a high frequency of *cpsY* mutations was observed among our LNZ-resistant strains (**Fig. S4**). Despite the prevalence of mutations in this LysR-family transcriptional regulator^44^, there is currently no direct evidence linking such mutations to LNZ resistance.

In addition to distinct mutations in primary resistance loci, secondary mutations were common across our resistant strains, and no two strains shared the same combination of primary and secondary mutations. To evaluate the potential role of the observed mutations in resistance or collateral responses, we applied partial least squares regression to assess if specific loci were associated with changes in susceptibility to antibiotics involved in the conserved and potentially exploitable CS pairs we identified (FUS, LNZ, and RIF). As expected, the analysis assigned high weight to mutations in the primary resistance loci, reinforcing their strong association with increased MICs for their corresponding antibiotics. Given their close association with each resistance phenotype, these primary resistance loci also emerged as potential contributors to the observed CS effects. However, whether mutations in these loci alone are sufficient to drive CS phenotypes remains inconclusive. Our analysis also identified a few secondary loci whose mutations may be associated with decreased MICs, suggesting a possible link to CS (**Fig. 3**). A notable example of such a secondary mutation was found in *pde1* of all CIP- and FUS-resistant strains and in some LNZ- and RIF-resistant strains (**Fig. S2**). While previous studies observed that pde1 affects penicillin resistance, genome stability and osmoregulation^45–47^, its phenotypic impact on the susceptibility of the collaterally-affected antibiotic could not be determined in our study due to frequent co-occurrence with primary resistance mutations. Since many of these secondary mutations emerged alongside the primary resistance mutations and mostly occurred in no more than 5 mutant strains (**Fig. 3**), it remains difficult to determine their individual contribution to CS phenotypes, and investigating this further is beyond the scope of our current study.

**Fig. 3.**
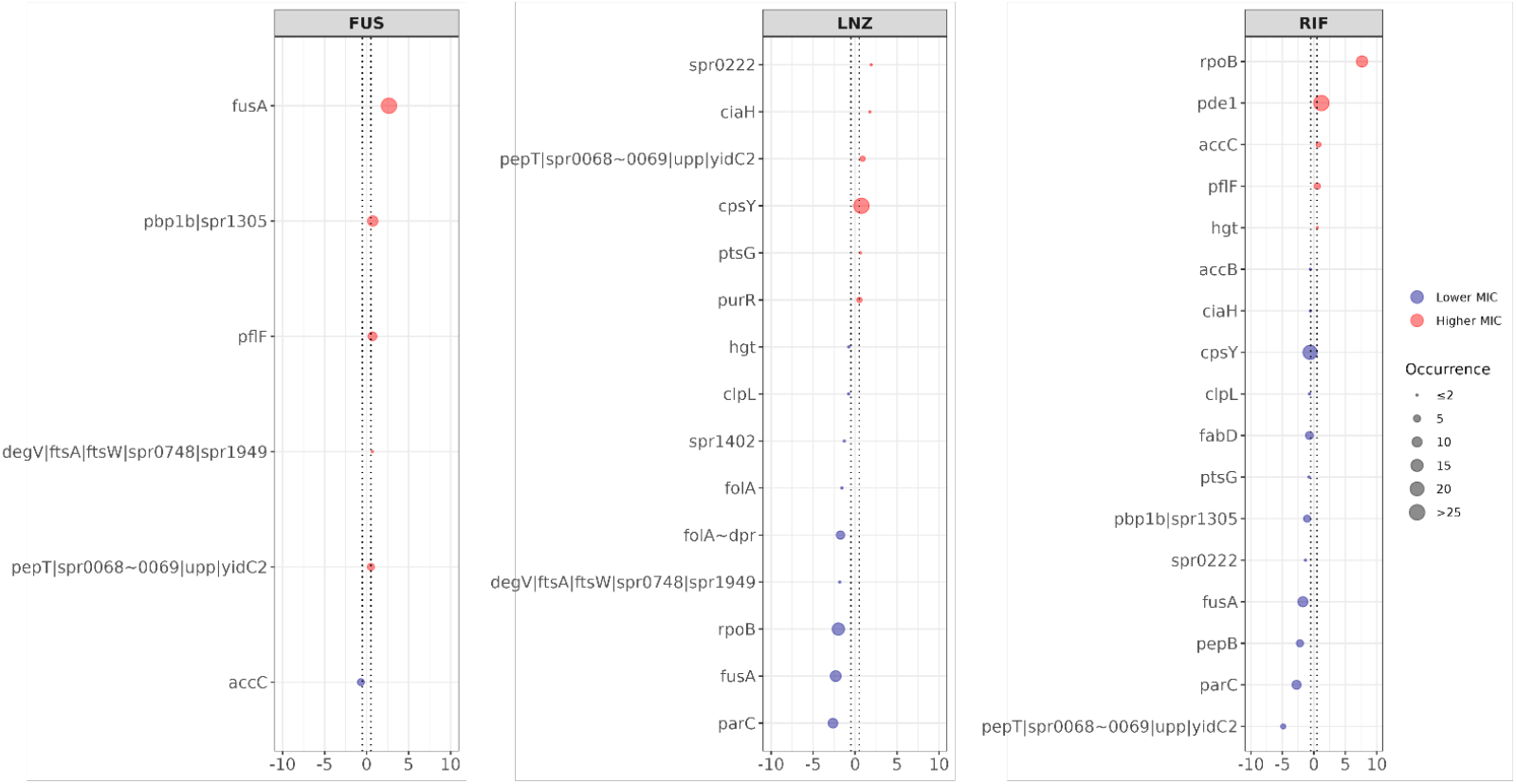
Impact of mutated loci or their combination on antibiotic susceptibility to FUS, LNZ, and RIF. The y-axis shows each mutated locus (or loci combinations), and the x-axis shows its effect on the log_2_-MIC ratio relative to the parental strain, quantified using partial least squares regression for each antibiotic. Positive coefficients indicate higher MICs (reduced susceptibility), and negative coefficients indicate lower MICs (increased susceptibility). An arbitrary cut-off of ±0.5 (dotted lines) was used to highlight loci or combinations with a stronger influence on MIC; those with estimated effects below this threshold were excluded from the figure. Colour indicates the direction of MIC change (red: increase, blue: decrease), while dot size reflects the frequency of the locus or combination among 50 resistant strains. Combinations are denoted by “ | “ between gene names, while mutations in the intergenic region between two genes are denoted by “ **∼** “.

### Identification of potentially exploitable CS antibiotic pairs for treatment

To identify antibiotic pairs that are potentially exploitable for the effective treatment of *S. pneumoniae* infections, we screened for pairs of antibiotics that consistently exhibited unidirectional or bidirectional CS without any evidence of CR. Eight unidirectional CS interactions were identified that appeared consistently in at least 9 out of 10 resistant strains (**Fig. 4A**). Among these eight antibiotic pairs displaying unidirectional CS, the RIF–FUS combination (RIF → FUS) appears particularly promising, as both FUS- and RIF-resistant strains exhibited high occurrence of CS alongside limited occurrence of CR, especially in the RIF-resistant mutants (**Fig. 2)**. Bidirectional CS pairs were rare and only emerged when the CS consistency threshold was reduced to 50% (CS_50_; **Fig. 4B**). Only two such interactions were observed: FUS–LNZ, where 5 out of 10 FUS-resistant mutants and 7 out of 10 LNZ-resistant mutants showed CS toward the other antibiotic (LNZ ↔ FUS), and FUS–SXT where 5 out of 10 FUS-resistant mutants and all 10 SXT-resistant mutants showed CS toward the other antibiotic (SXT ↔ FUS). Of the two, the pair FUS–LNZ appears more promising for the treatment of *S. pneumoniae* infections. This is based on the observation that SXT-resistant strains exhibited a high occurrence of CR, particularly towards CIP, CLI, GEN, and LNZ (**Fig. 2; bottom-left panel**), making the involvement of SXT resistance a less desirable option for clinical application. Additionally, SXT already consists of a combination of two antibiotics, making the pharmacodynamics of further combinations with FUS more complex. Given these considerations, we decided to quantitatively explore FUS–RIF and FUS–LNZ as suitable antibiotic pairs for CS-based treatment strategies.

**Fig. 4.**
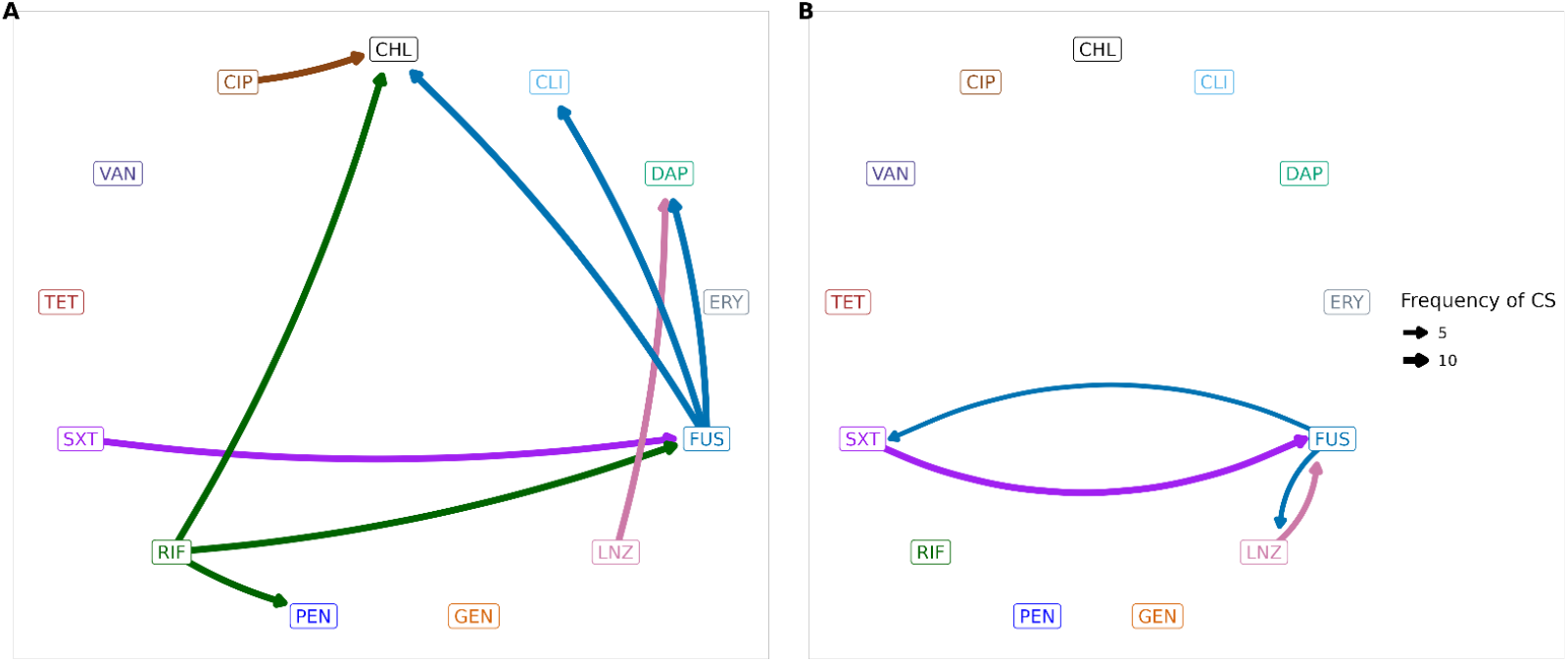
Conserved collateral sensitivity network. Each conserved collateral effect between two antibiotics is represented by an arrow. Arrows originate from the antibiotic to which the strain was resistant, with colour indicating that antibiotic, and point toward the antibiotic for which a collateral sensitivity effect was observed. Arrow width indicates the frequency of the collateral sensitivity among the 10 mutant strains resistant to each respective antibiotic. (A) Unidirectional CS interactions observed in at least 9 out of 10 resistant strains. (B) Bidirectional CS interactions observed in at least 5 out of 10 resistant strains.

### CS-informed combinations suppress the *de novo* establishment of resistant bacterial populations

Having identified the FUS–RIF and FUS–LNZ as potentially promising antibiotic pairs for CS-informed therapies, we first performed checkerboard assays, which indicated no physiological interaction between these antibiotic pairs (FICI = 2 for both). We then conducted static time-kill experiments to derive the growth rate as well as the PD characteristics of FUS, LNZ, and RIF on the parental strain (**Table 1, Fig. S5**). The relationship between antibiotic concentration and bactericidal effect was described using the pharmacodynamic parameters EC_50_ (the concentration producing 50% of the maximum effect), Emax (the maximum bactericidal effect), and the Hill coefficient (which quantifies the steepness of the concentration– effect curve). The bactericidal activity of RIF demonstrated a markedly steeper concentration–effect relationship, with a Hill exponent of 6.57 (arbitrary units), indicating a sharp transition between sub-therapeutic and effective concentrations. In contrast, the effects of FUS and LNZ were adequately described by a standard Emax model with the Hill exponent fixed at 1.00, reflecting a more gradual dose–response relationship.

**Table 1.**
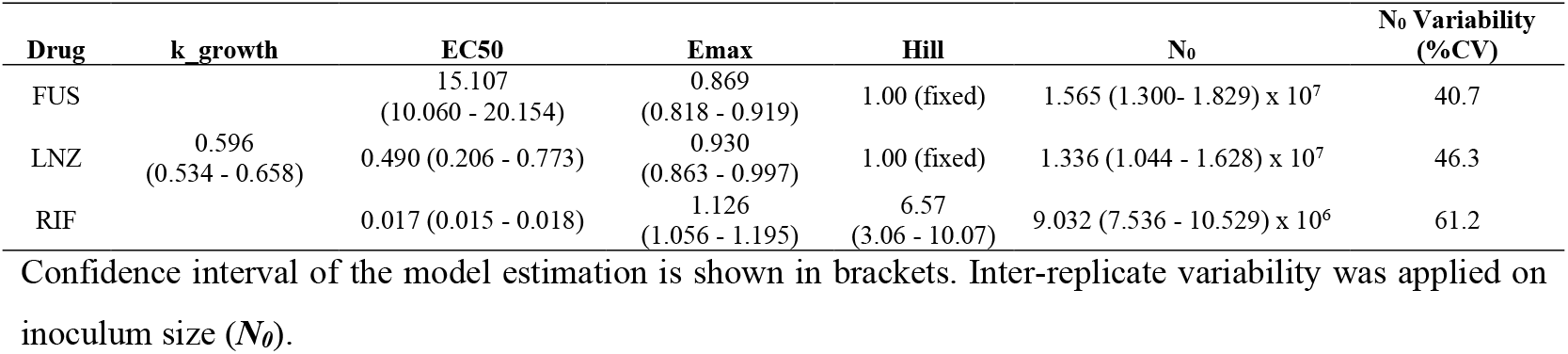
Estimated PD Characteristics of FUS, LNZ, and RIF on *S. pneumoniae* R6.

| Drug | k<sub>growth</sub> | EC<sub>50</sub> | Emax | Hill | N<sub>0</sub> | N<sub>0</sub> Variability (%CV) |
| --- | --- | --- | --- | --- | --- | --- |
| FUS |  | 15.107<br>(10.060 - 20.154) | 0.869<br>(0.818 - 0.919) | 1.00 (fixed) | 1.565 (1.300- 1.829) x 10<sup>7</sup> | 40.7 |
| LNZ | 0.596<br>(0.534 - 0.658) | 0.490 (0.206 - 0.773) | 0.930<br>(0.863 - 0.997) | 1.00 (fixed) | 1.336 (1.044 - 1.628) x 10<sup>7</sup> | 46.3 |
| RIF |  | 0.017 (0.015 - 0.018) | 1.126<br>(1.056 - 1.195) | 6.57<br>(3.06 - 10.07) | 9.032 (7.536 - 10.529) x 10<sup>6</sup> | 61.2 |

Confidence interval of the model estimation is shown in brackets. Inter-replicate variability was applied on inoculum size (*N*<sub>0</sub>).

Next, we used these estimated PD parameters (**Table 1**) in subsequent simulations to evaluate the outcomes of antibiotic treatments involving FUS, LNZ, and RIF, administered either as monotherapy or in pairwise combinations. Our monotherapy simulations showed that resistance rapidly evolved in all or most simulations. Specifically, the standard clinical FUS dosing regimen (500 mg three times daily) was not effective in suppressing infection. Instead, a FUS-resistant population was predicted to become established within 4 days (**Fig. 5A-B**), likely due to the high plasma protein binding of FUS^32^, which reduces the free (active) FUS concentration in the plasma and results in a low overall FUS exposure. The probability of suppressing the infection and achieving a 10-fold reduction in bacterial density using standard LNZ (600 mg twice daily) and RIF (300 mg twice daily) monotherapies did not exceed 8.4% and 28.2%, respectively (**Fig. 5A**). LNZ- and RIF-resistant populations were similarly predicted to become established within 4 days during treatment (**Fig. 5C-D**). Our results suggest that standard monotherapy with FUS, RIF, or LNZ is unlikely to effectively suppress bacterial growth or prevent resistance establishment during treatment.

**Fig. 5.**
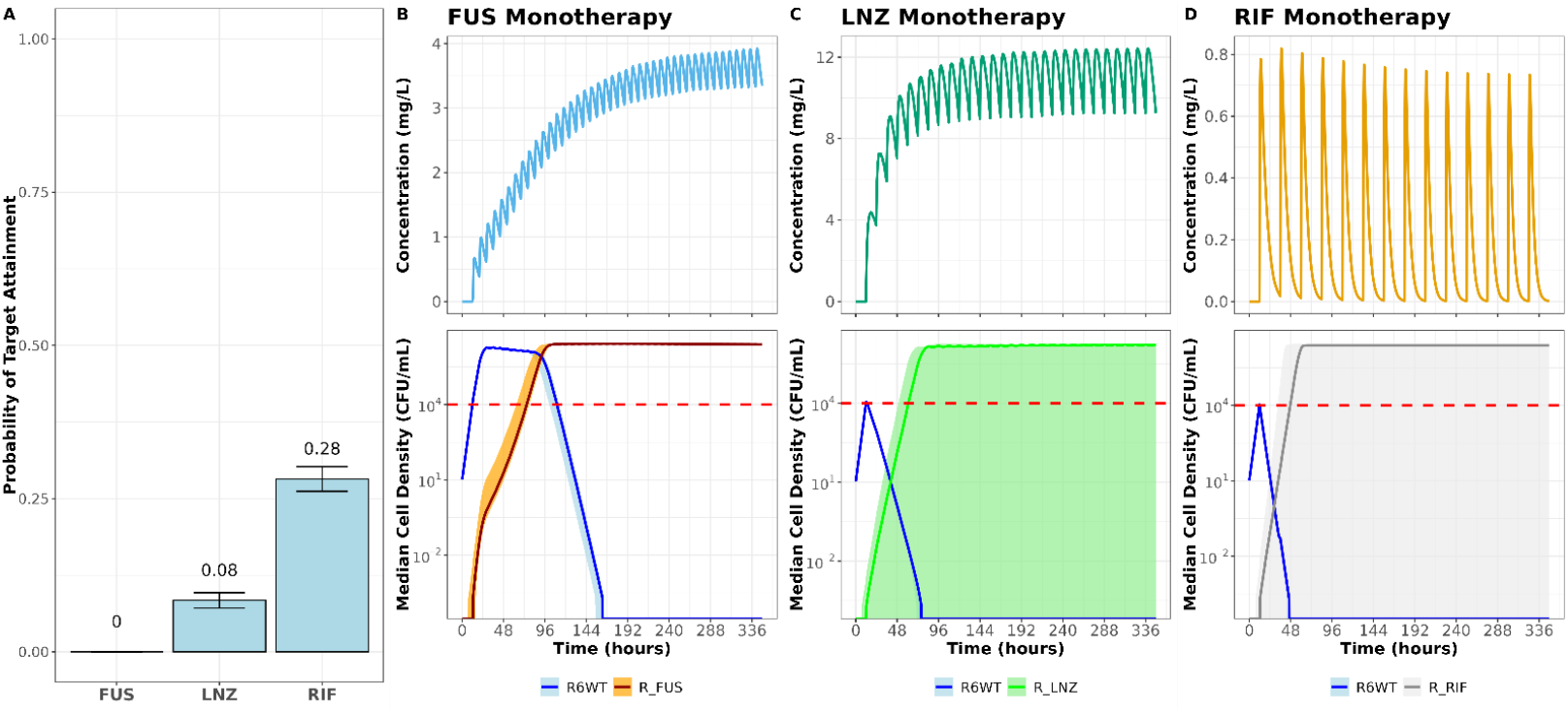
Effectivity of clinical monotherapy treatment of FUS, LNZ, and RIF. **A)** Probability of suppressing bacterial density to below 10^3^ CFU/mL at the end of treatment with FUS, LNZ, and RIF monotherapies. Error bars represent the standard error of the probability of treatment success. **B–D)** Time-course profiles of standard monotherapy with FUS, LNZ, and RIF, respectively. Solid lines represent the median of the simulations, and shaded areas indicate the 95% confidence interval (2.5th to 97.5th percentile). The horizontal dashed red line indicates the bacterial density at the start of treatment (10^4^ CFU/mL). Sub-populations resistant to an antibiotic not used in each monotherapy, arbitrarily chosen from the other two antibiotics, were included in the simulations to ensure an equivalent overall resistance rate across all scenarios, but are not shown in the plot, as their density remained below 1 CFU/mL in all cases.

To assess the efficacy of CS-based combination therapy with FUS-LNZ and FUS-RIF we tested three dosing schedules: (1) co-administration (mixing) with a 24-hour dosing interval, and sequential administration with 12-hour dose-cycling intervals starting with either (2) FUS or, depending on the combination, (3) either LNZ or RIF. The FUS-LNZ pair exhibiting fairly conserved bidirectional CS (FUS ↔ LNZ) was highly effective in suppressing *S. pneumoniae* infection. This combination therapy was able to achieve the target level of bacterial suppression with greater than 95% probability (**Fig. 6A**). Moreover, resistance to either antibiotic was not predicted to emerge during treatment with this combination (**Fig. 6B-D**). The success of the combination was relatively insensitive to the dosing schedule used. However, the effectiveness of the combination depended highly on CS interactions between FUS and LNZ. In the absence of CS, the highest achievable probability of target attainment of the combination was reduced to 68.0%. The FUS-RIF pair exhibiting conserved unidirectional CS (RIF → FUS) also showed improved target attainment compared to the respective monotherapies. However, its overall efficacy was substantially lower than that of the FUS-LNZ combination, with a maximum attainment rate of only 39.8%. Notably, co-administration was more effective than the other tested dosing strategies, while sequential administration starting with FUS was the least effective. This suggests that rapidly suppressing the bacterial population at the onset of treatment is critical for this combination (**Fig. 6A**) and confirms the importance of drug order in cases of unidirectional CS. As with the FUS-LNZ combination, the efficacy of FUS-RIF also appeared to depend on the CS interaction between the two antibiotics, but to a lesser degree (**Fig. 6A**). To evaluate how dependent the success of the CS-based combination therapy is to the consistency of observed CS among the 10 tested strains for each antibiotic resistant group, additional dosing schedules were tested where we removed the collateral effects from 2, 5, or 8 strains with the highest CS magnitude. This revealed that for FUS-LNZ, even the collateral effect removal from just the top two strains lead to an approximately 10% drop in the target attainment. In contrast, FUS-RIF showed a more gradual decline in efficacy, indicating a broader but weaker reliance on CS consistency across strains (**Supplementary Material 2**). Although combining the antibiotics offered some improvement over monotherapy, the emergence of a resistant population remained likely with the FUS-RIF combination (**Fig. 6E-G**). Results from our simulations highlight that CS-informed antibiotic combinations are more effective at eradicating bacterial infections and suppressing the *de novo* emergence of resistance, but their efficacy depends on the directionality and consistency of CS and on the dosing schedule.

**Fig. 6.**
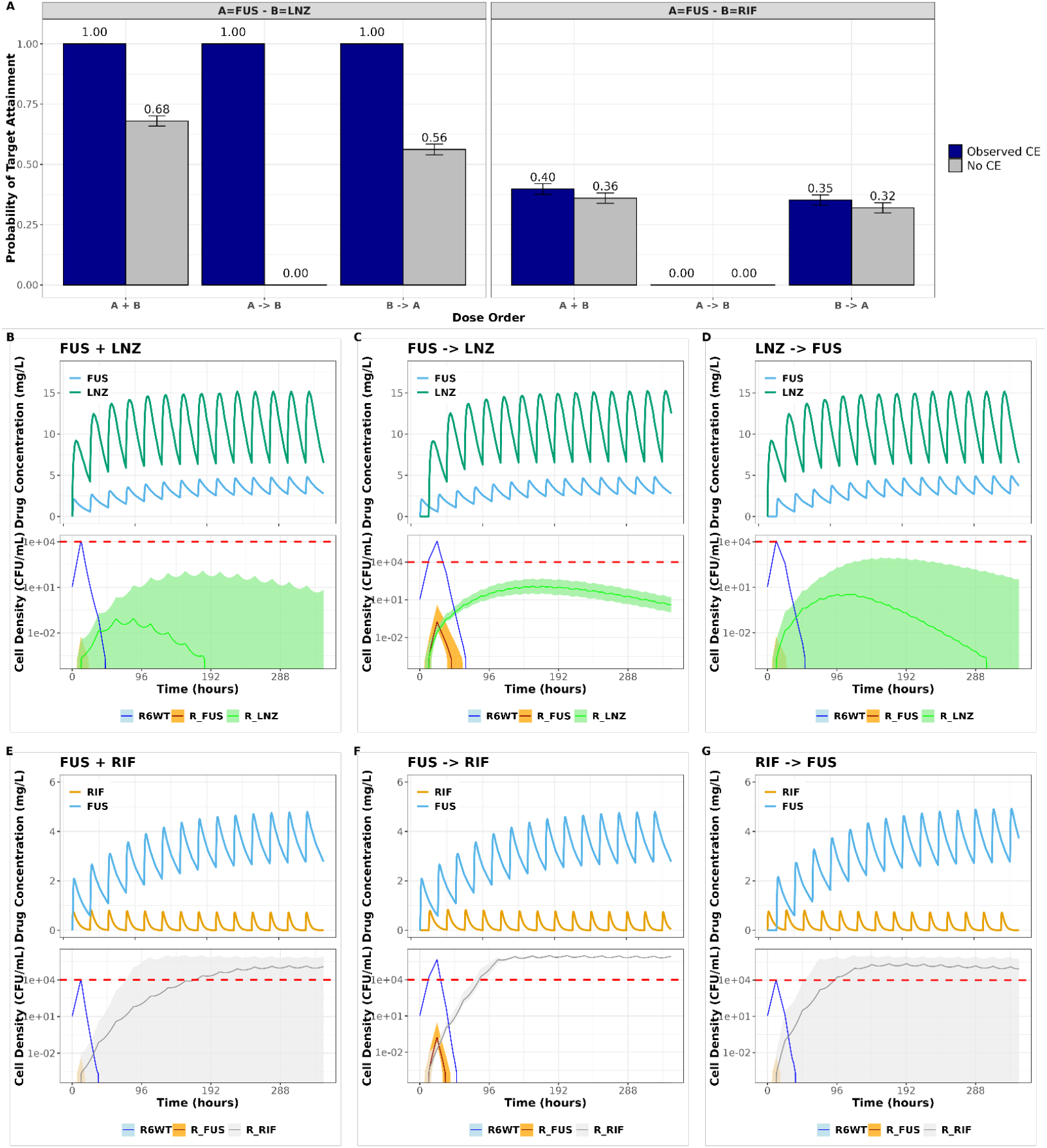
Efficacy of CS-based FUS-LNZ and FUS-RIF combination. **A)** Probability of suppressing bacterial density to below 10^3^ CFU/mL at the end of treatment. Three modes of dose administration time were evaluated, namely co-administration (A+B), 12-hour cycling starting with antibiotic A (A→B), and 12-hour cycling starting with antibiotic B (B→A). Treatment outcome with the observed and without collateral effects (CE) were evaluated. The error bars represent the standard error of the probability of treatment success. **B–G)** Time-course profiles of CS-based FUS-LNZ and FUS-RIF combination therapies. Solid lines represent the median of the simulations, and shaded areas indicate the 95% confidence interval (2.5th to 97.5th percentile). The horizontal dashed red line indicates the bacterial density at the start of treatment (10^4^ CFU/mL).

## Discussion

CS offers a promising strategy to combat antibiotic resistance by exploiting trade-offs that arise during the evolution of resistance to existing antibiotics^11,13^. However, the clinical utility of CS-based approaches depends on the consistency of CS phenotypes across independent resistance mutants. This is particularly relevant for pathogenic bacteria such as *S. pneumoniae*, in which rapid within-host evolution has been documented, sometimes resulting in antibiotic resistant genotypes that can later become established during antibiotic treatment^48,49^. Although *S. pneumoniae* is included in the priority list of bacterial pathogens in need of effective treatment options^50^ and ranks among the top five pathogens globally for deaths attributable to and associated with antibiotic resistance^51^, current knowledge of collateral sensitivity in this species remains limited, having been investigated only in the context of a single antibiotic class^12^. To address this gap and capture CS effects associated with early and stochastic evolutionary events that give rise to resistant variants prior to antibiotic exposure, events that may occur even within hosts and seed resistance under clinical conditions, we employed a fluctuation-based approach. This allowed us to recover multiple spontaneous resistant mutants against antibiotics with distinct modes of action commonly used against Gram-positive infections and to systematically identify robust CS interactions in *S. pneumoniae*.

We identified a high number of CS interactions in our study, suggesting that *de novo* resistance to certain antibiotics can reliably produce collateral vulnerabilities. Moreover, consistent CS was observed despite genotypic heterogeneity among our mutant strains, indicating that such vulnerabilities can arise even if there are divergent mutational pathways towards resistance. Among the resistance phenotypes tested, RIF-resistance induced the most consistent CS across the majority of the antibiotics, which aligns with earlier observations in *Escherichia coli*^52^. By contrast, RIF-resistance in *Enterococcus faecalis* conferred extensive but heterogenous collateral effects depending on the antibiotic^53^. Our findings also highlight interspecies conservation of several unilateral CS interactions between *S. pneumoniae* and *E. coli* (RIF → VAN)^52^, *S. pneumoniae* and *E. faecalis* (RIF → CHL)^53^, *S. pneumoniae* and *Enterococcus faecium* (RIF → CLI and RIF → ERY)^54^ and *S. pneumoniae* and *Staphylococcus aureus* (FUS → ERY)^55^, indicating that CS-based antibiotic combinations may have generalizable therapeutic potential across diverse bacterial species.

Despite growing interest in CS, which has led to the identification of CS interactions across various bacterial pathogens, the underlying mechanisms of these interactions remain largely unresolved for most interactions^9^. To explore the genetic basis of CS in *S. pneumoniae*, we performed whole-genome sequencing and applied partial least squares regression to identify specific loci and/or mutations associated with collateral responses. However, directly attributing CS to individual genomic changes proved challenging as mutations in known resistance-associated loci frequently co-occurred with secondary mutations, complicating causal interpretation. Notably, no single mutation was found universally across all resistant mutants, indicating that the consistent CS observed likely resulted from multiple distinct mutational patterns that converged on similar phenotypic outcomes.

Nonetheless, several of the mutations that we identified in known resistance-associated loci have previously been shown to exert pleiotropic effects on bacterial physiology, which may be sufficient to drive the observed CS responses. For example, mutations in the β-subunit of RNA polymerase (*rpoB*)^52,56^ and in topoisomerase IV (*parC*)^57^ have been shown to alter transcription and DNA supercoiling, respectively, leading to global effects on gene expression. Mutations in elongation factor G (*fusA*)^58,59^ have been associated with reduced levels of the stress sensor ppGpp and increased sensitivity to oxidative stress and DNA damage. Alterations in dihydrofolate reductase (*folA*)^60^ affect tetrahydrofolate biosynthesis, a precursor for thymidylate and purine production, and have been linked to widespread shifts in the bacterial proteome and metabolome. Likewise, linezolid resistance mutations, such as those in 23S rRNA, ribosomal proteins, or RNA methyltransferases^61^ have been shown to impair ribosomal function and reduce protein synthesis efficiency. While these pleiotropic effects suggest a potential mechanistic basis for the consistent CS in *S. pneumoniae*, the presence of co-occurring secondary mutations in our strains prevents definitive conclusions. Nevertheless, the associations identified here provide a valuable starting point for future studies, involving targeted gene editing and phenotypic testing in isogenic strains to determine whether these mutations play a direct role in mediating CS.

Our experimental findings demonstrate that similar CS can arise through distinct evolutionary paths, reinforcing its potential robustness as a therapeutic strategy. To further evaluate this potential, we integrated our experimental data with a pharmacokinetics and pharmacodynamics model simulating the development and treatment of *S. pneumoniae* bacteraemia. For this, we focused on an antibiotic pair exhibiting unidirectional CS (RIF → FUS) and a pair showing bidirectional CS (LNZ ↔ FUS). These CS-based combinations were evaluated *in silico* under simultaneous administration and 12-hours cycling regimens starting with either antibiotic compared to monotherapy, showing that therapies based on such antibiotic combinations can be more effective than monotherapy in limiting the emergence of resistance and eliminating the infection. Among the combinations tested, the bidirectional CS observed between FUS and LNZ appeared to offer greater therapeutic benefit compared to the unidirectional CS observed between FUS and RIF. Importantly, the order of antibiotic administration influenced treatment outcomes. Starting with the antibiotic whose resistance leads to the strongest CS effect, such as LNZ or RIF in this study, was consistently more effective in constraining resistance and eliminating the infection. While co-administration showed the greatest potential in suppressing mutant establishment, previous studies have raised concerns about the risk of selecting double-resistant mutants under simultaneous antibiotic pressure^62^. This possibility was not directly evaluated in the current study, but future work should investigate whether co-administration promotes dual resistance and how such outcomes might impact the long-term success of CS-based combination therapies.

Interestingly, the identification of a CS-based combination involving LNZ is particularly significant given the role of LNZ as a critical last-resort antibiotic for the treatment of severe infections caused by Gram-positive pathogens, including multidrug-resistant *S. pneumoniae*^63,64^. The ability to pair LNZ with a collateral-sensitizing partner such as FUS may provide a valuable strategy to enhance its efficacy while simultaneously minimizing the risk of resistance development. This is especially relevant in light of recent reports indicating a rising prevalence of LNZ-resistance among *S. pneumoniae* clinical isolates^65^, which threatens to compromise its long-term therapeutic utility. Thus, incorporating CS-based strategies could contribute to extending the clinical lifespan of LNZ by mitigating selective pressure that drives resistance evolution.

In conclusion, our study demonstrates the potential of collateral sensitivity to inform more effective antibiotic combination strategies against *S. pneumoniae*. By integrating experimental evolution, genomic analysis, and computational modelling, we identified reproducible CS interactions and evaluated their utility in guiding treatment design. While the genetic basis of CS remains complex and requires further investigation, particularly regarding the role of secondary mutations, our findings support the feasibility of leveraging CS to prevent the establishment of resistance, even within heterogeneous bacterial populations. By exploring the consistency of CS development in *S. pneumoniae* and evaluating its potential therapeutic value, this study lays the foundation for future research into the clinical applicability of CS-based treatment strategies aimed at preserving antibiotic efficacy and limiting resistance establishment during *S. pneumoniae* infections and their treatment.

## Data availability

All data needed to evaluate the conclusions of this paper are present in the paper and/or the supplementary material accompanying it. Raw data are available at DataverseNL (xxxxxx). Whole-genome sequencing data are available at the National Center for Biotechnology Information (BioProject PRJNA224116). The model and the associated code are available at Github (xxxxxx)

## Contributions

A.L. and D.E.R. designed the study; A.L., S.T.T., E.F., and I.V.H. performed experiments and acquired the data; A.L., S.T.T., and B. V. performed the data analysis; A.L. and S.T.T. prepared the first draft of the manuscript; all authors interpreted the data, read, contributed in the writing of, and approved the final manuscript.

