## Supplementary material for "Divergent spontaneous antibiotic-resistance evolution confers reciprocal and exploitable collateral sensitivity effects"

### Supplementary Figures

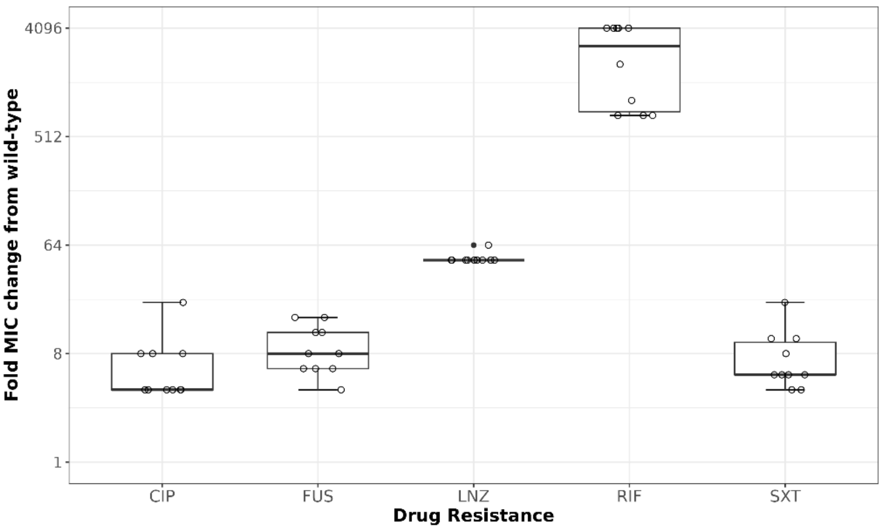

**Fig. S1 | Fold change in MICs of antibiotic-resistant mutant strains.** The fold change in MICs was determined for ten antibiotic-resistant strains against ciprofloxacin (CIP), fusidic acid (FUS), linezolid (LNZ), rifampicin (RIF), and trimethoprim-sulfamethoxazole (SXT). Values are shown as fold changes relative to the wild-type R6 strain. Boxes indicate the interquartile range (IQR) with the median line, whiskers extend to 1.5×IQR, and closed circles (●) represent outliers beyond 1.5×IQR. Open circles (○) represent individual MIC measurements for each resistant strain.

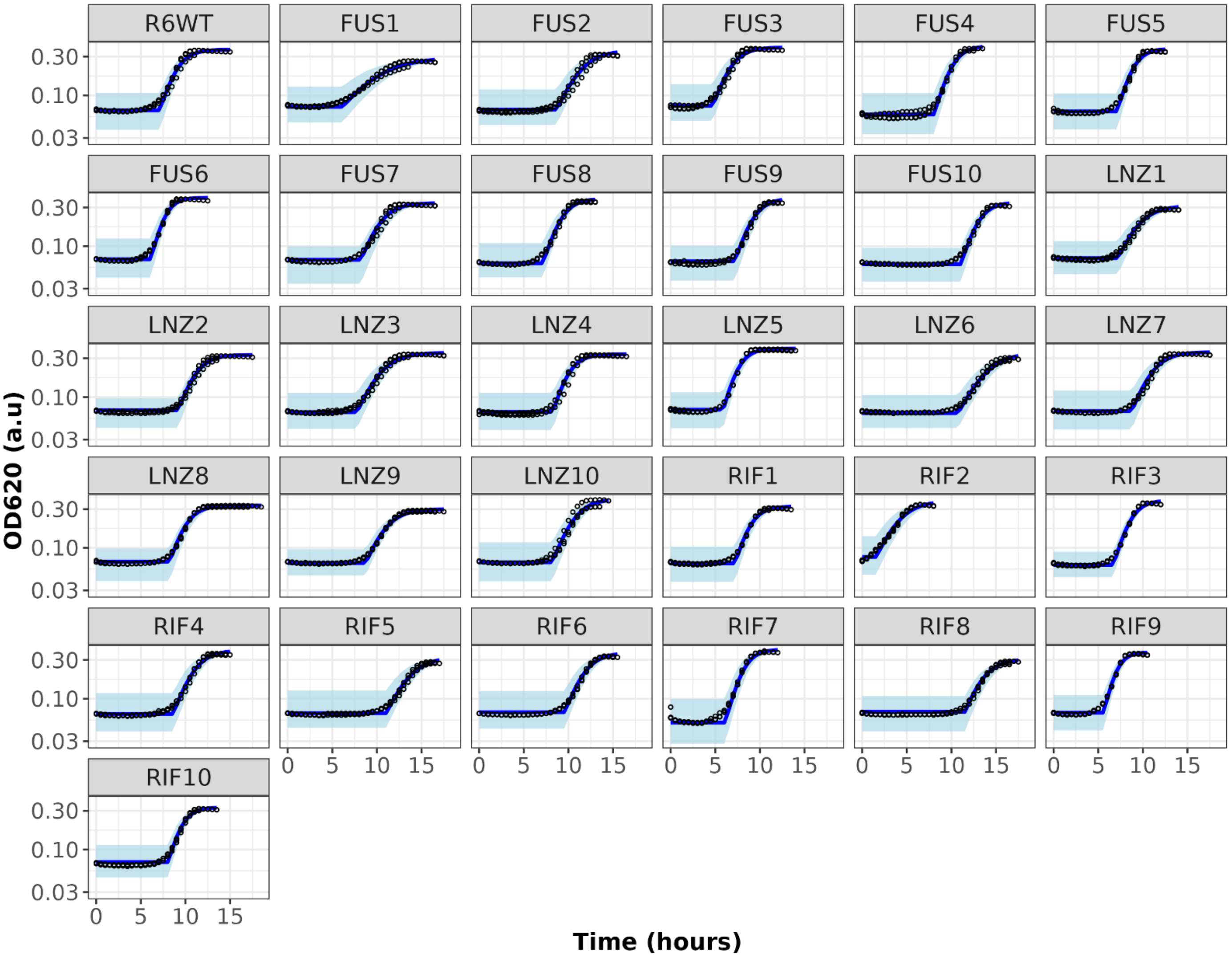

**Fig. S2 | Characterization of parental and mutant *S. pneumoniae* strains.** Growth of the bacterial culture was monitored based on changes in OD620. The growth rate and maximum cell density were estimated from three observation replicates. Dots represent each experimental observation, while the solid blue line represents the median prediction of the strain growth by the model. The shaded area represents the 95% prediction interval of the model.

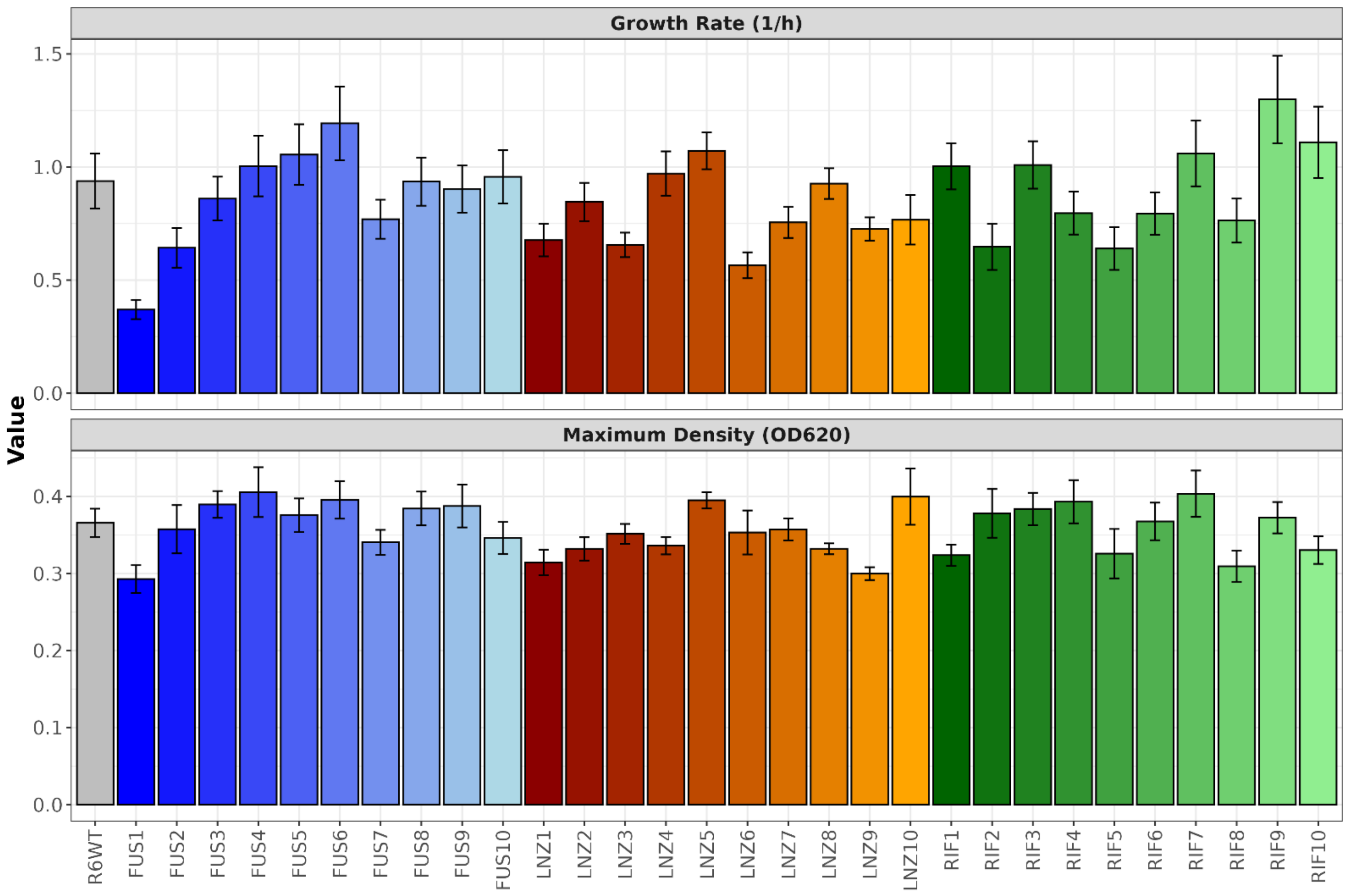

**Fig. S3 | Growth characteristics of parental and mutant strains.** Growth characteristics were summarized by the estimated growth rate ($h^{-1}$) and maximum cell density ($OD_{620}$; a. u.) parameters. Height of the bar and the error bars represented the estimate, and the 95% confidence interval of the parameter fit, respectively. Parameter fit was derived from three observation replicates. Mutants belonging to the same drug-resistance group are depicted using the same color gradient, with mutants resistant to FUS shown in blue, to LNZ in green, and to RIF in red.

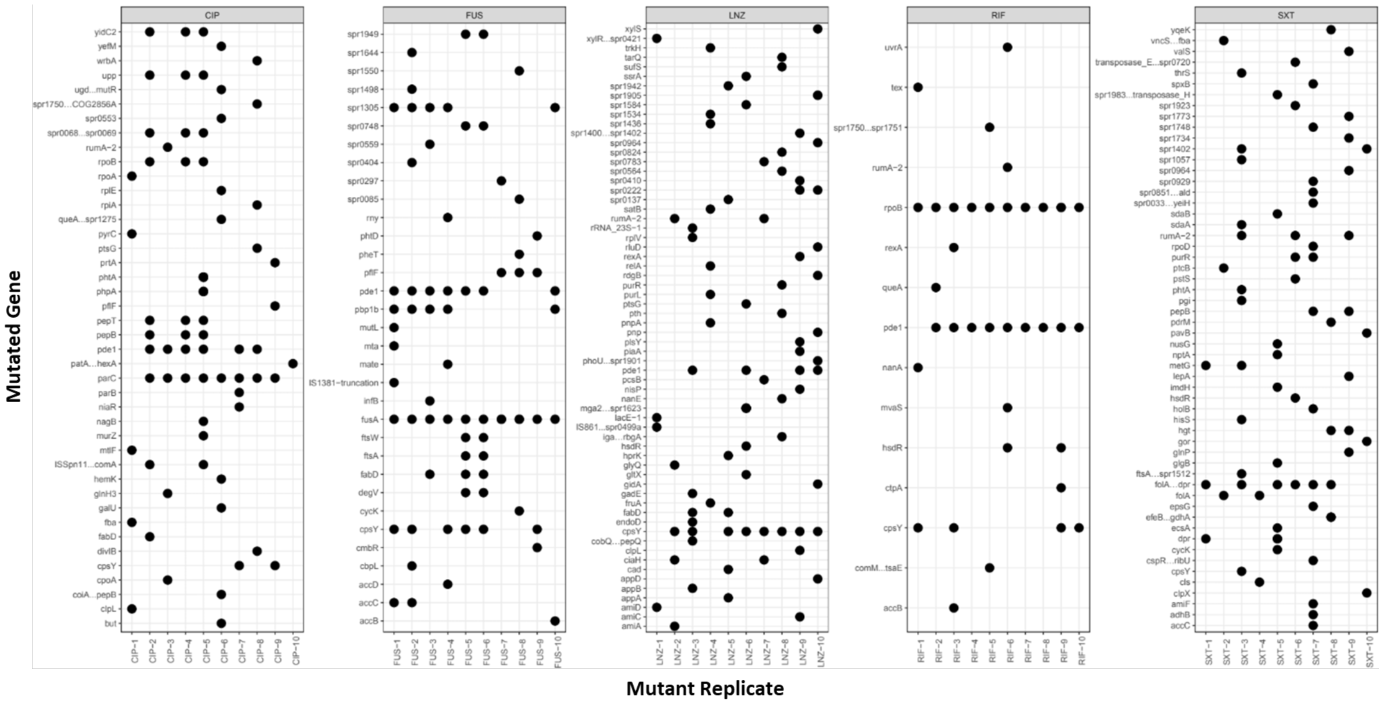

**Fig. S4 | Mutational patterns in antibiotic-resistant strains.** Mutations in 185 loci were identified across 50 antibiotic-resistant strains. The x-axis represents mutant replicates, while the y-axis denotes the mutated gene identifiers. Each point indicates the presence of a mutation in a specific gene within a given strain. Overall, LNZ- and SXT-resistant mutants exhibited a broader range of affected genes, whereas RIF-resistant strains displayed a more limited set of mutations.

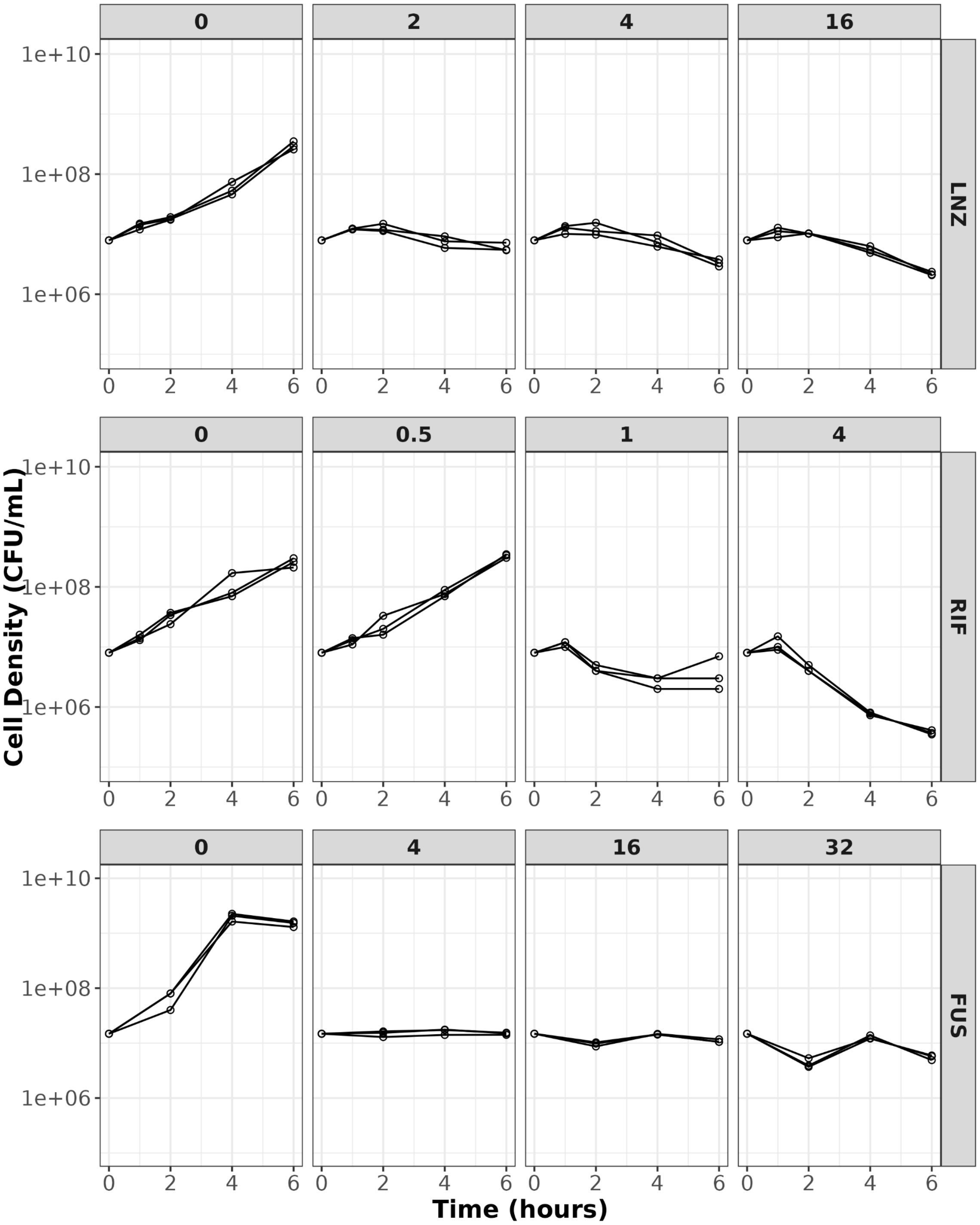

**Fig. S5 | Growth curve of strain R6 under static FUS, LNZ, and RIF concentrations.** Static time-kill experiments were conducted over 6 hours, with each condition tested in triplicate. Individual measurements are shown as points, and consecutive measurements from the same replicate are connected by straight lines.

### Supplementary Tables

**Table S1.** ***Streptococcus pneumoniae*** **strains included in this study and their characteristics.**

| **Strains** | **Relevant characteristic(s)** | **Source** |
|---|---|---|
| R6 WT | Pan-susceptible, non-encapsulated strain derived from the serotype 2 *Streptococcus pneumoniae* D39 clinical strain | Laboratory stock |
| ATCC49619 | Encapsulated strain of serotype 19F, quality control for susceptibility testing | Laboratory stock |
| CIP-1, CIP-2, CIP-3, CIP-4, CIP-5, CIP-6, CIP-7, CIP-8, CIP-9, CIP-10 | Fluoroquinolone-resistant strain derived from *Streptococcus pneumoniae* R6 WT | This study |
| FUS-1, FUS-2, FUS-3, FUS-4, FUS-5, FUS-6, FUS-7, FUS-8, FUS-9, FUS-10 | Fusidic acid-resistant strain derived from *Streptococcus pneumoniae* R6 WT | This study |
| LNZ-1, LNZ-2, LNZ-3, LNZ-4, LNZ-5, LNZ-6, LNZ-7, LNZ-8, LNZ-9, LNZ-10 | Oxazolidinone-resistant strain derived from *Streptococcus pneumoniae* R6 WT | This study |
| RIF-1, RIF-2, RIF-3, RIF-4, RIF-5, RIF-6, RIF-7, RIF-8, RIF-9, RIF-10 | Rifampicin-resistant strain derived from *Streptococcus pneumoniae* R6 WT | This study |
| SXT-1, SXT-2, SXT-3, SXT-4, SXT-5, SXT-6, SXT-7, SXT-8, SXT-9, SXT-10 | Trimethoprim/sulfamethoxazole-resistant strain derived from *Streptococcus pneumoniae* R6 WT | This study |

**Table S2. List of antibiotics used in this study and their targets.**

| Antibiotic Name (Abbreviation) | Antibiotic class | Antibiotic target(s) | Supplier |
|---|---|---|---|
| Chloramphenicol (CHL) | Amphenicol | Protein synthesis (50S) | Sigma-Aldrich |
| Ciprofloxacin (CIP) | Quinolone | DNA replication (GyrA + ParC) | Acros Organics |
| Clindamycin (CLI) | Lincosamide | Protein synthesis (50S) | Cayman chemical |
| Daptomycin (DAP) | Lipopeptide | Cell membrane | Acros Organics |
| Erythromycin (ERY) | Macrolide | Protein synthesis (50S) | Sigma-Aldrich |
| Fusidic Acid (FUS) | Fusidane | Protein synthesis (EF-G) | Cayman chemical |
| Gentamicin (GEN) | Aminoglycosides | Protein synthesis (30S) | Sigma-Aldrich |
| Linezolid (LNZ) | Oxazolidinone | Protein synthesis (50S) | Cayman chemical |
| Penicillin (PEN) | β-lactam | Cell wall synthesis (PBPs) | Sigma-Aldrich |
| Rifampicin (RIF) | Rifamycin | RNA synthesis (rpoB) | GERBU Biotechnik |
| Trimethoprim/ sulfamethoxazole (SXT) | Antifolate | Folate synthesis (FolA + FolP) | SERVA/ AG Scientific |
| Tetracycline (TET) | Tetracycline | Protein synthesis (30S) | Sigma-Aldrich |
| Vancomycin (VAN) | Glycopeptide | Cell wall synthesis | Carl Roth |

**Table S3. MIC of *S. pneumoniae* R6 WT**

| Antibiotics | MIC (mg/L) |
|---|---|
| CIP | 0.7500 |
| FUS | 8.0000 |
| LNZ | 1.0000 |
| RIF | 0.0195 |
| SXT | 0.1875 |
| CHL | 3.0000 |
| CLI | 0.0625 |
| DAP | 0.0938 |
| ERY | 64.0000 |
| GEN | 1.5000 |
| PEN | 0.0234 |
| TET | 0.1875 |
| VAN | 0.0938 |

**Table S4. Individual characteristics used in PK models.**

| Characteristics | Value | Units | Notes |
|---|---|---|---|
| *Bodyweight* | 70 | kg | Used as standard value in FUS and LNZ models[1, 2]. |
| *Age* | 69 | years | Median age reported in the LNZ reference. Age was only used in LNZ PK model[2]. |
| *Creatinine clearance* | 6 | L/h-70kg | Used as standard value in LNZ reference. Creatinine clearance was only used in LNZ PK model[2]. |

**Table S5. Standard doses of FUS, LNZ, and RIF in the Netherlands.**

| Drug | Dose (mg) | Administration | Reference |
|---|---|---|---|
| FUS | 500 (1500 mg total daily) | three times daily – oral dose | 7 |
| LNZ | 600 (1200 mg total daily) | twice daily – oral dose | 8 |
| RIF | 600 (600 mg total daily) | once daily – oral dose | 9 |

**Table 6. Growth characteristics of mutant and parental *S. pneumoniae* strains**

| Strain | $k_{growth}$ | $N_{max}$ | $t_{delay}$ | $N_0$ | $N_0$ Variability (%CV) |
|---|---|---|---|---|---|
| R6WT | 0.938 (0.816 - 1.06) | 0.366 (0.348 - 0.384) | 7.13 (6.977 - 7.283) | 0.067 (0.065 - 0.07) | 30 |
| FUS1 | 0.37 (0.327 - 0.412) | 0.293 (0.275 - 0.311) | 6.127 (5.912 - 6.342) | 0.075 (0.074 - 0.077) | 25 |
| FUS2 | 0.643 (0.555 - 0.731) | 0.357 (0.326 - 0.389) | 8.671 (8.49 - 8.852) | 0.065 (0.062 - 0.069) | 29.5 |
| FUS3 | 0.86 (0.763 - 0.957) | 0.39 (0.372 - 0.407) | 4.733 (4.596 - 4.87) | 0.075 (0.071 - 0.079) | 28.6 |
| FUS4 | 1.004 (0.87 - 1.138) | 0.406 (0.373 - 0.438) | 8.068 (7.927 - 8.208) | 0.058 (0.054 - 0.061) | 28.7 |
| FUS5 | 1.055 (0.921 - 1.189) | 0.376 (0.354 - 0.397) | 7.129 (7.001 - 7.257) | 0.063 (0.062 - 0.065) | 27.9 |
| FUS6 | 1.193 (1.03 - 1.356) | 0.396 (0.371 - 0.42) | 6.131 (6.007 - 6.256) | 0.07 (0.068 - 0.072) | 28.1 |
| FUS7 | 0.769 (0.683 - 0.856) | 0.34 (0.324 - 0.357) | 8.213 (8.066 - 8.359) | 0.067 (0.065 - 0.068) | 28.6 |
| FUS8 | 0.935 (0.829 - 1.042) | 0.384 (0.362 - 0.406) | 7.154 (7.028 - 7.28) | 0.063 (0.061 - 0.064) | 27.6 |
| FUS9 | 0.903 (0.798 - 1.007) | 0.388 (0.36 - 0.415) | 7.193 (7.075 - 7.312) | 0.063 (0.062 - 0.064) | 26.5 |
| FUS10 | 0.956 (0.839 - 1.074) | 0.346 (0.325 - 0.367) | 11.071 (10.944 - 11.199) | 0.062 (0.061 - 0.063) | 26 |
| LNZ1 | 0.677 (0.605 - 0.749) | 0.314 (0.298 - 0.331) | 7.206 (7.077 - 7.335) | 0.07 (0.068 - 0.073) | 24.3 |
| LNZ2 | 0.845 (0.761 - 0.93) | 0.332 (0.317 - 0.347) | 9.183 (9.067 - 9.298) | 0.066 (0.064 - 0.069) | 25.4 |
| LNZ3 | 0.656 (0.601 - 0.71) | 0.351 (0.338 - 0.364) | 7.762 (7.635 - 7.889) | 0.066 (0.064 - 0.068) | 26.2 |
| LNZ4 | 0.971 (0.873 - 1.069) | 0.336 (0.325 - 0.347) | 8.254 (8.141 - 8.367) | 0.064 (0.06 - 0.068) | 27.6 |
| LNZ5 | 1.071 (0.99 - 1.152) | 0.395 (0.384 - 0.406) | 5.883 (5.805 - 5.962) | 0.068 (0.067 - 0.07) | 25.5 |
| LNZ6 | 0.565 (0.508 - 0.623) | 0.353 (0.325 - 0.382) | 10.759 (10.634 - 10.884) | 0.066 (0.065 - 0.066) | 23.8 |
| LNZ7 | 0.755 (0.686 - 0.824) | 0.357 (0.343 - 0.372) | 8.754 (8.631 - 8.877) | 0.066 (0.065 - 0.068) | 26.9 |
| LNZ8 | 0.927 (0.859 - 0.994) | 0.332 (0.325 - 0.339) | 8.243 (8.155 - 8.33) | 0.066 (0.065 - 0.067) | 23.6 |
| LNZ9 | 0.726 (0.675 - 0.777) | 0.3 (0.292 - 0.308) | 8.749 (8.658 - 8.84) | 0.066 (0.065 - 0.067) | 21.8 |
| LNZ10 | 0.767 (0.657 - 0.876) | 0.4 (0.363 - 0.436) | 8.197 (8.031 - 8.364) | 0.067 (0.064 - 0.07) | 30.5 |
| RIF1 | 1.003 (0.901 - 1.106) | 0.324 (0.31 - 0.338) | 7.217 (7.118 - 7.316) | 0.066 (0.065 - 0.067) | 24.6 |
| RIF2 | 0.647 (0.545 - 0.749) | 0.378 (0.346 - 0.41) | 1.576 (1.358 - 1.794) | 0.079 (0.076 - 0.083) | 28.9 |
| RIF3 | 1.009 (0.904 - 1.114) | 0.384 (0.363 - 0.405) | 6.709 (6.606 - 6.813) | 0.063 (0.061 - 0.064) | 26 |
| RIF4 | 0.796 (0.701 - 0.89) | 0.393 (0.365 - 0.421) | 8.622 (8.48 - 8.764) | 0.065 (0.064 - 0.066) | 27.5 |
| RIF5 | 0.64 (0.545 - 0.734) | 0.326 (0.294 - 0.358) | 11.051 (10.88 - 11.221) | 0.066 (0.065 - 0.067) | 25.8 |
| RIF6 | 0.794 (0.7 - 0.888) | 0.368 (0.343 - 0.392) | 9.592 (9.455 - 9.728) | 0.066 (0.065 - 0.067) | 26 |
| RIF7 | 1.06 (0.915 - 1.205) | 0.404 (0.373 - 0.434) | 6.106 (5.958 - 6.253) | 0.055 (0.054 - 0.057) | 31 |
| RIF8 | 0.763 (0.666 - 0.86) | 0.309 (0.289 - 0.329) | 11.51 (11.367 - 11.653) | 0.066 (0.065 - 0.067) | 24.1 |
| RIF9 | 1.299 (1.106 - 1.492) | 0.372 (0.352 - 0.393) | 5.619 (5.491 - 5.746) | 0.068 (0.066 - 0.07) | 28.3 |
| RIF10 | 1.109 (0.951 - 1.267) | 0.33 (0.312 - 0.348) | 8.079 (7.945 - 8.212) | 0.067 (0.066 - 0.068) | 26.8 |
